# Dynamic mean and variance of microparasite load give key insights into population dynamics and underlying mechanisms

**DOI:** 10.1101/2024.09.24.614578

**Authors:** Jason Cosens Walsman, Sabrina H Streipert, Cheryl J Briggs, Mark Q Wilber

**Affiliations:** University of California Santa Barbara; University of Pittsburgh; University of Tennessee Knoxville

**Keywords:** Epidemiology, within-host dynamics, trait heterogeneity, parasite aggregation, host regulation

## Abstract

Heterogeneity among individuals, in number of parasites, body size, etc., can have critical interactions with population dynamics. We tease these out with a relatively simple model of such interactions in a case study of microparasite load with empirically supported assumptions to ask a key question. How does variance in microparasite load interact with population-level dynamics? We show how the mean and variance of infection load vary throughout an epidemic. Further, we show how mean and variance have mutual negative feedbacks on each other mediated by high disease-induced death rates at high loads. Helpfully, we find that mean and variance provide information into underlying processes as well. Population-level trends in the mean and variance reveal underlying trends in within-host processes, e.g., differentiating host evolution of defence that manifests as tolerance, constitutive resistance, inducible resistance, or acquired resistance. Such inference may be critical for managing endangered Mountain Yellow Legged Frog populations recovering from *Batrachochytrium dendrobatidis* epidemics. Lastly, we demonstrate the impact of pathogen load variance on host fitness, pathogen fitness, and host population depression by pathogens. Our results demonstrate the importance of trait heterogeneity and the insights available from relatively simple models, both for microparasite infection load and possibly other traits.

## Introduction

Individual organisms differ from each other critically in traits such as body size, the number of parasites they host, and vital rates with implications for population dynamics including evolution, ecological resilience, and our ability to make ecological inference. Quantitative genetics models have long considered how the mean and variance (most often small and fixed but not always) of heritable traits impact population dynamics [1,2]. Non-heritable sources of trait heterogeneity, our focus here, may also be of critical importance. Individual-level heterogeneity can interact with population dynamics for traits such as parasite load [3], body size [4], or vital rates [5]. Further, heterogeneity interacts with evolution as heterogeneity may slow evolution [6] or selection may favor alleles that alter variance in fitness, as classically expressed in terms of geometric mean fitness [7]. Further, heterogeneity can interact with ecological resilience, reflecting population declines or helping drive recoveries [4]. Additionally, trends in heterogeneity may contain important information about unobserved biological processes [8]. For these reasons, ecologists have frequently sought to understand and model individual heterogeneity in populations.

But one barrier to understanding the role of heterogeneity is the complexity of jointly modelling heterogeneity and population dynamics, particularly in such a way as to gain general analytical insight. For example, Partial Differential Equations, Integral Projection Models, Class-structured Matrix Models (for discrete traits) and Individual Based Models all provide approaches to relatively easily simulate the interaction between individual heterogeneity and population dynamics, but going beyond simulation-based results often requires biologically unrealistic assumptions or advanced mathematical techniques. In general, the literature is lacking somewhat for simpler models that i) allow for the joint modelling of individual heterogeneity and population dynamics ii) are based on biologically and empirically supported assumptions and iii) provide novel and generalizable conceptual insight.

Such models are particularly needed in epidemiology where individual heterogeneity among hosts can have substantial feedbacks on host and parasite population dynamics. For example, one major aspect of individual heterogeneity in host-parasite systems is that the minority of the hosts hold the majority of parasites. Such variance (or aggregation) in the number of parasites on each infected host (infection load) holds implications for host population dynamics [3,9,10] and variance patterns in wild populations can indicate within-host mechanisms [8,11]. These observations highlight the need for model-based, conceptual insight to address our focal question. How does variance in microparasite load interact with population-level dynamics? Classically, variance in infection load has been considered important for macroparasites (e.g., arthropods or helminths) but often ignored for microparasites (e.g., fungi, bacteria, viruses). In fact, microparasitic infections have often been treated as present or absent [12]. This binary view reflects an assumption that variation in microparasite infection load is negligible for one of two reasons; either, load variation is negligible because it is small due to homogeneous hosts and rapid microparasite growth to a steady state or load variation need not be small but lacks impact on key outcomes.

But many microparasite taxa display significant variance in infection load with important impacts on population dynamics, evolution, and resilience ecology. For example, HIV patients display wide variability in viral load, impacting HIV spread and likely also viral evolution [13]. Extreme variation in SARS-CoV-2 viral load may be critical for superspreading [14]. Variation in bacterial pathogen load can help predict clinical outcomes for patients [15,16] and likely also impacts pathogen evolution as bacterial genotypes vary in their loads [16]. The fungal agent of White Nose Syndrome achieves very different pathogen loads in bats, both across hibernation duration, within species at a given time, and between bat species with conservation implications [17]. Frogs infected with the fungal agent of chytridiomycosis, *Batrachochytrium dendrobatidis* (*Bd*), display widely varying infection loads within and between populations, impacting the survival of individual frogs as well as the persistence of populations [18]. Because heterogeneity in microparasite load is critical for many taxa, we need new theory for how this heterogeneity interplays with population dynamics.

While existing theory for macroparasite load provides a critical foundation, we need further development for microparasites. Microparasite load, as quantified by molecular techniques (e.g., qPCR), can follow continuous distributions of infection load [14,19–21] rather than the discrete distributions of macroparasite counts. Further, the availability of laboratory data allows us to parameterize more biologically realistic, non-linear functions for the relationships between infection load and key processes such as pathogen growth rate or the mortality of infected hosts [22] contrasted with the linear functions typically used in classic macroparasite theory. Seizing these opportunities, we develop novel theory for heterogeneity in microparasite load focused on dynamic mean and variance in load. With a simple model (relative to models with full, arbitrary trait distributions like Integral Projection Models), we address our focal question of how microparasite load variance interacts with population dynamics. We show how the mean and variance i) are affected by population dynamics throughout epidemic stages ii) affect population dynamics, especially host suppression and iii) have mutual negative feedbacks on each other leading to multiple possible qualitative patterns for load distributions. Importantly, one implication of these cross-scale, bi-directional feedbacks is that the dynamic patterns of mean and variance in infection intensity contain distinct, qualitative signatures of biological mechanisms that are impossible to identify from data such as trends in abundance. We illustrate this phenomenon with a case study of recovering populations of endangered Mountain Yellow Legged Frogs depressed by *Bd*. Using our model, we demonstrate that the mean and standard deviation of infection load provide enough information to infer the type of host defence strategy that hosts are evolving to recover following disease-induced declines, differentiating tolerance, constitutive resistance, inducible resistance, and acquired resistance. Altogether, we find that a relatively simple model of heterogeneity gives us new insights into population dynamics, evolution, and our ability to infer within-individual processes from population-level patterns.

### Model structure

Trait heterogeneity within a population interacts with the population-level processes, birth, death, infection, etc., that determine how many individuals are in the population. We model these interactions with an Integral Projection Model (IPM) that tracks the population density of susceptible hosts, infected hosts, and environmental pathogens (“zoospores” per our focal case study of amphibian chytrid) at each time step (*t, t*+1, etc). We use one day time steps, but this assumption is easily relaxed [23]. We assume that log load *x* is normally distributed in the class of infected hosts, a reasonable assumption for *Bd* [24] and many other microparasites [13,14,20,21]. This assumption allows us to derive a much more tractable and simpler model (see Appendix for derivation of our reduced dimension IPM from the full IPM via moment closure), in which we only need to track the mean and standard deviation of log load rather than the full distribution. This simpler model allows us to analytically tease out how the mean and variance of log load impact population dynamics and vice versa (Fig. 1). This is all but impossible to do with the full IPM directly. Three key processes govern load dynamics. First, newly infected hosts tend to have low loads. Second, pathogen growth increases loads. Third, infected hosts with higher loads are more likely to die.

**Figure 1.**
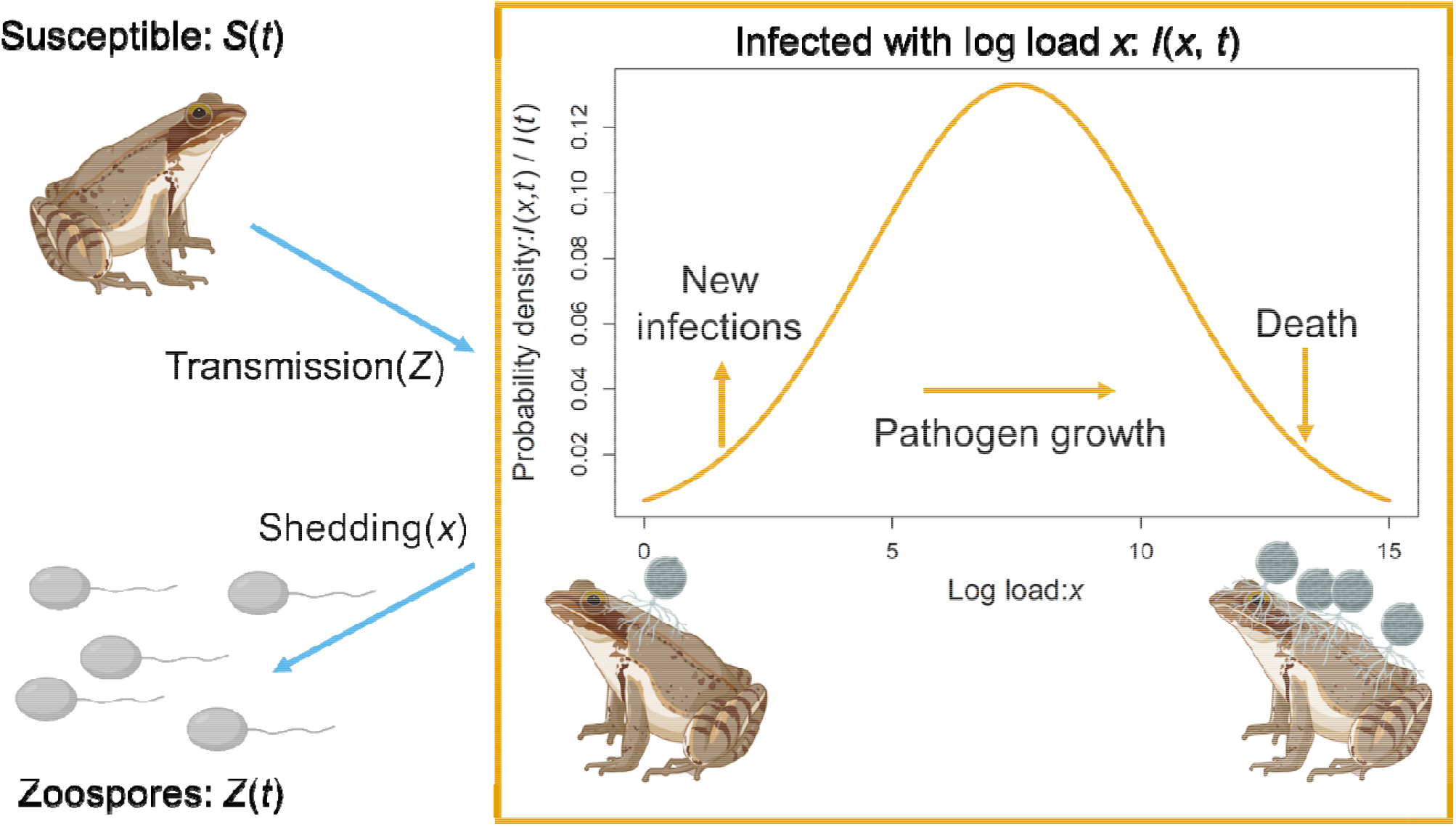
Conceptual diagram of disease model. Hosts begin life susceptible then transmission moves them into the infected class, which is structured by the distribution of log pathogen load, *x*, at a given time, *t*. New infections predominantly add hosts at low loads, bringing the mean load down. Pathogen growth increases loads, increasing the mean. Loss of infection, partially from host recovery but predominantly from host death, disproportionately occurs at high loads and brings the mean down. Pathogens are shed back into the environment at a rate proportional to load. *I*(*x, t*) d*x* is the population density of hosts with a load between *x* and *x* + d*x* at time *t*, and *I*(*t*) is the total population density of infected hosts at time *t*.

Further, we consider host evolution in this model, for our focal case study of recovering frog populations, via changes in genotype frequencies. For simplicity, we assume clonal competition among frog genotypes where those that are more defended in terms of their epidemiological traits have lower fecundity (*r*_*j*_); in particular, we assume this defence-fecundity trade-off has a curvature that favors evolution toward intermediate values of defence in the presence of pathogens. In each time step, susceptible hosts of genotype *j* are born when susceptible or infected hosts [*S*_*j*_(*t*)+*I*_*j*_(*t*)=*N*_*j*_(*t*)] reproduce, with some negative density dependence (*e*^-*qN*(*t*)^ where *N*(*t*) is all hosts and *q* is the strength of negative density dependence), and some probability of mutation (*m*, Eq. 1a). Negative density dependence that regulates host populations in the absence of disease.

Susceptible hosts may survive (probability *s*_0_) and remain uninfected [depending on transmission rate *β*_*j*_ and zoospore density *Z*(*t*)] or become infected while infected hosts that survive may recover back to the susceptible class (with probability *l*, Eq. 1b and 1c). The probability an infected host survives a time step declines with log load *x* in a non-linear fashion [survival = 1-□((*x*-*μ*_LD50, *j*_)/*σ*_LD50_) where □ is the cumulative distribution function of the standard normal, an empirically supported functional form [22]]; higher values of *σ*_LD50_ model a more gradual, sigmoidal decline of survival probability from 1 to 0 and *μ*_LD50,*j*_ gives the LD50 of load. The average survival probability of infected hosts of genotype *j* always declines with their mean load, *μ*_*j*_(*t*) [average survival: *ŝ*(*α*_j_(*t*)) where *ŝ*(*z*) = 1-□(*z*) and 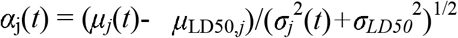 is a quantity that arises multiple times]. Higher standard deviation of load, *σ*_*j*_(*t*), leads to higher death rates, as long as the minority of infected hosts are dying each time step, because a higher standard deviation increases the minority of hosts with high and deadly loads [∂*ŝ*(*α*_j_(*t*))/∂*σ*_*j*_(*t*)<0 if and only if *μ*_*j*_(*t*) < *μ*_LD50_, i.e., *ŝ*(*α*_j_(*t*)) > 0.5].

The dynamics of the mean and variance are captured in the dynamics of the first and second moment of log load. Mean log load is related to the first moment of log load [*μ*_*j*_(*t*) = *P*_*j*_(*t*)/*I*_*j*_(*t*); *P*_*j*_(*t*) = ∫*xI*_*j*_(*x,t*)d*x* and *I*_*j*_(*t*) = ∫*I*_*j*_(*x, t*) d*x* over all *x* where *I*_*j*_(*x,t*)d*x* is the population density of hosts of genotype j with a log infection load between *x* and *x* + d*x*]; the first moment captures all sources that contribute to pathogen load (Eq. 1d) as newly infected hosts have low log loads (drawn from a normal distribution with mean *μ*_*0*_ and standard deviation *σ*_*0*_) and pathogens grow within hosts with some variability and negative density dependence [*x*(*t*+1) is drawn from a normal distribution with mean *a+bx*(*t*) and standard deviation *σ*_G_; 0<*b*<1 captures negative density dependence]; this functional form of microparasite growth is empirically supported for our focal system [22] and such within-host growth on a log scale up to a carrying capacity may be appropriate for many host-microparasite systems. The second moment of log load is related to the standard deviation of log load [*σ*_*j*_(*t*) = (*Q*_*j*_(*t*)/*I*_*j*_(*t*) - *μ*_*j*_^2^(*t*))^1/2^ where *Q*_*j*_(*t*) = ∫*x*^2^*I*_*j*_(*x,t*)d*x* over all *x*]; this second moment captures all sources that contribute to log load and variation in log load such as variation in the loads of newly infected hosts and variation in pathogen growth (e.g., unlike Eq. 1d, *σ*_0_ and *σ*_G_ appear in Eq. 1e). Key to both the first and second moments of log load (and thus mean and standard deviation) are the three processes controlling pathogen load: new infections, pathogen growth, and death (see Fig. 1). Both the mean and standard deviation of log load contribute to the mean of load on the natural scale (*e*^*μ*+*σ*^2/2^ is mean load on the natural scale) and thus pathogen shedding from infected hosts (with λ giving the zoospores shed per within-host pathogen per time unit) and pathogens persist for some time in the environment (with probability *v* per time step, Eq. 1f).

<u>Eq. 1a: Birthrate of genotype</u>*<u>j</u>*

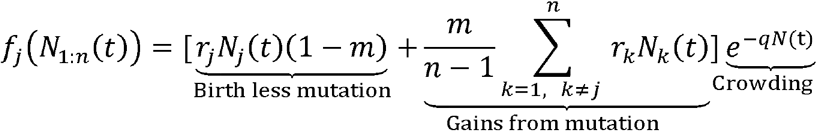

<u>Eq. 1b for susceptible individuals of genotype j</u>

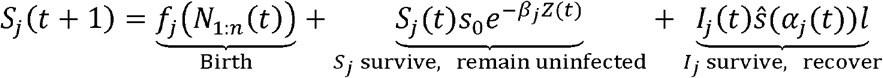

<u>Eq. 1c for infected individuals of genotype j</u>

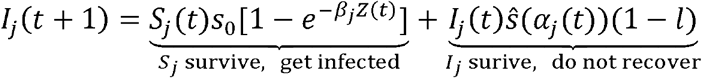

<u>Eq. 1d for first moment of log load on infected individuals of genotype j (“0^th^” etc. refer to terms of pathogen growth of different order with respect to pathogen density dependence,</u>*<u>b</u>*<u>_j_)</u>

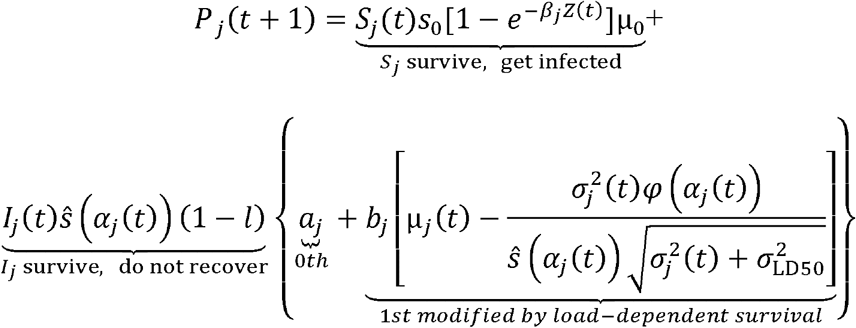

<u>Eq. 1e for second moment of log load on infected individuals of genotype j</u>

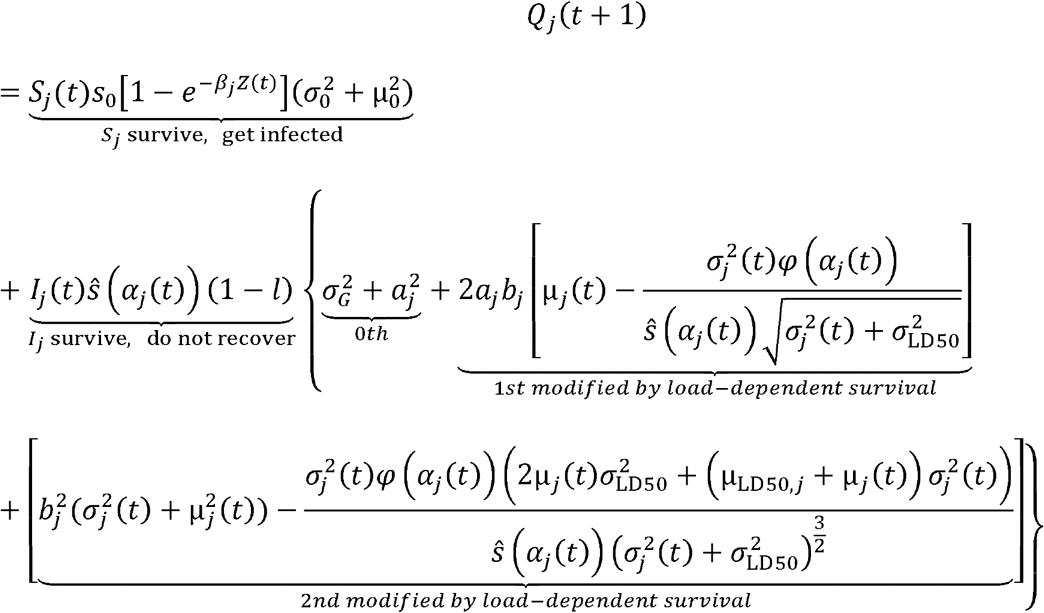

<u>Eq. 1f: Environmental pathogens</u>

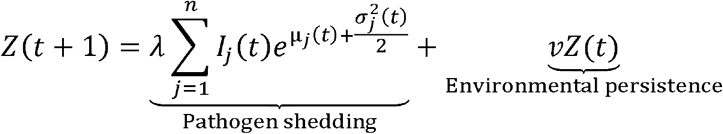

Note that *φ*(*z*) is the probability density function of the standard normal distribution at *z* [*φ*(*z*)=(1/√(2π)) *e*^-(*z*^2)/2^]. We also derive a very similar version of the model in which recovered hosts enter a recovered class with acquired immunity (until immunity wanes) such that they have different traits upon reinfection and the load of reinfected hosts follows its own lognormal distribution (see Appendix).

### Load dynamics from epidemic outbreak to endemic equilibrium

With these equations for the dynamics of the host population, pathogen population, and the distribution of load, we can gain insight into how population dynamics impact load. We simulated a small density of hosts who just became infected in a host population of one genotype at its disease-free equilibrium (using biologically reasonable parameters for Mountain Yellow Legged Frogs and *Bd*; see Appendix for values and simulation procedure). At different stages of the epidemic (beginning, log growth, peak, and endemic equilibrium), we see different distributions of load reflecting how the population-level processes impact the distribution of load (beginning, log growth, and peak stages are marked by coloured squares in Fig. 2A).

**Figure 2.**
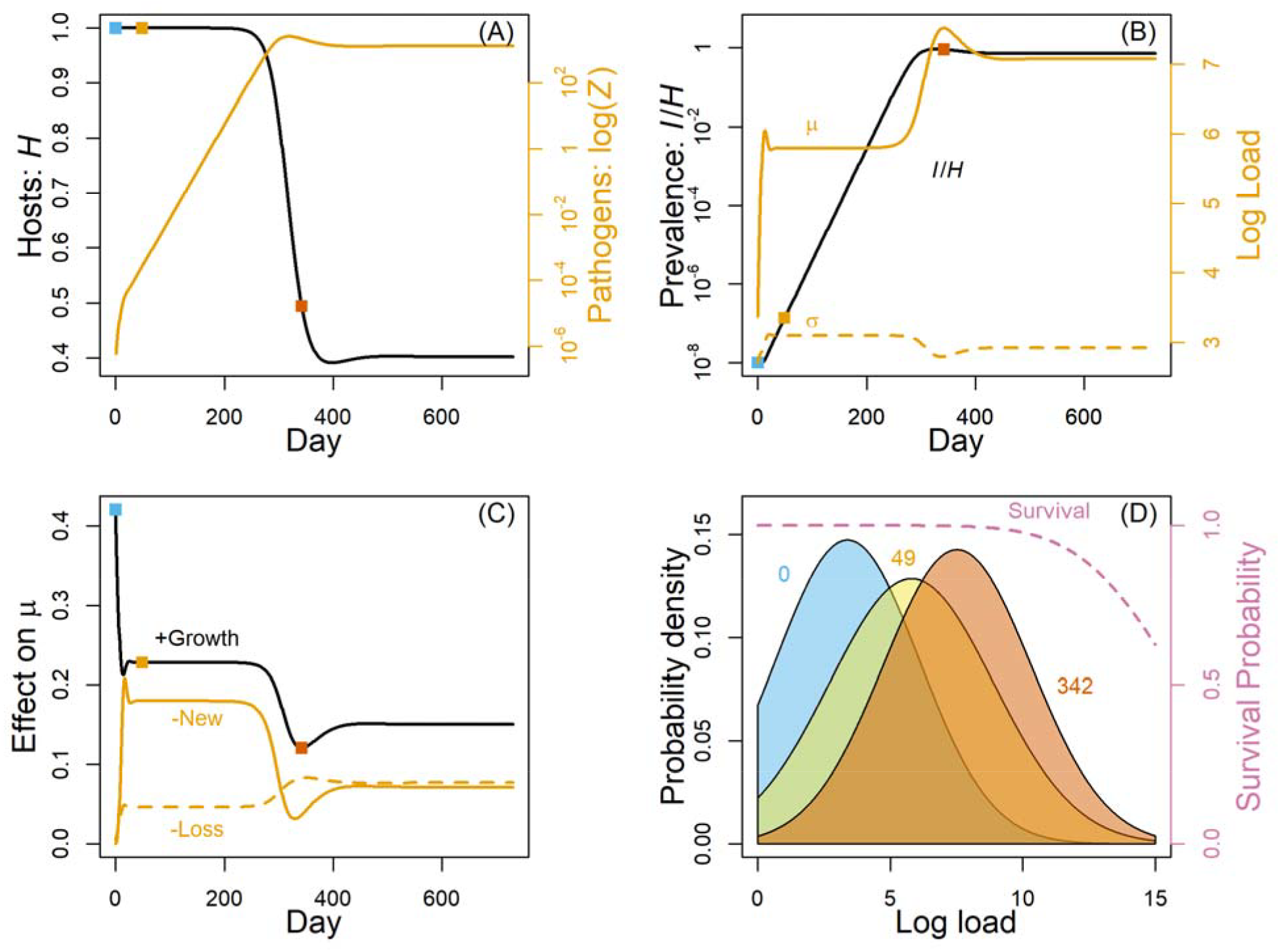
Epidemic stage shifts the distribution of pathogen load. (A) As a rare pathogen outbreaks, host density (black curve) declines and pathogen density (gold curve) rises. Coloured squares denote epidemic stages of beginning (blue), log growth (yellow), and peak (orange). (B) Prevalence (black) rises over time until it reaches a peak then settles slightly lower toward equilibrium. Mean (solid gold) and standard deviation (dashed gold) of log load change during these epidemic stages as well. (C) The change in the mean is determined by a positive contribution from pathogen growth (black) and negative contributions from new infections (solid gold) and infection loss (dashed gold), which is predominantly due to high load infections being lost at a higher rate. (D) The bell curve of log load shifts higher or lower and broader or narrower at different epidemic stages (days 0, 49, and 342 shown). Survival declines with log load, particularly at high loads, constraining the distribution of load (dashed pink).

At the beginning of the epidemic, there are very few infected hosts and those who are infected have only just become infected [i.e., *μ*(*t* = 0)=*μ*_0_ and *σ*(*t* = 0) = *σ*_0_], causing low mean and standard deviation of load (Day 0 in Fig. 2B). Pathogen growth during the log growth of prevalence stage (Day 49 in Fig. 2B) quickly raises the mean load and there is now separation in the infected population between newly infected hosts and hosts who have been infected long enough to experience pathogen growth (Fig. 2C shows how pathogen growth raises the mean while new infections bring it down) so there is now a wider spread of loads [i.e., *σ*(*t* = 49) > *σ*(*t* = 0), compare blue and yellow bell curves in Fig. 2D]. As the rate of increase of prevalence slows and even becomes slightly negative (from around day 200 to orange square in Fig. 2B), mean log load rises and the standard deviation declines (compare orange curve for peak stage to yellow log growth curve in Fig. 2D). Mean load rises again at this stage because there are far fewer new, low load infections entering the infected class and dragging the mean down (see sharp decrease in New curve in Fig. 2C). The higher mean pushes the right tail of the load distribution up against the survival constraint (Fig. 2D, orange bell curve’s right tail extends far into the declining portion of the pink survival curve); loss of these high load infections shrinks the standard deviation of load down [see how *σ*(*t*) has declined at day 342 in Fig. 2B]. Loads have become so high that the survival constraint forces mean loads back down to approach their endemic equilibrium value and the standard deviation is allowed to recover to somewhat higher values [see rising Loss curve in Fig. 2C around day 342 as well as decreasing *μ*(*t*) and increasing *σ*(*t*) in Fig. 2B from day 342 onward]. These general patterns for *μ*(*t*) and *σ*(*t*) hold across a broad range of parameters (see code), capturing the impacts of epidemic stage on load.

The impact of epidemic stage on the distribution of load also prompts investigation of the long-term, endemic equilibrium mean and standard deviation. We found implicit expressions for the equilibrium mean (*μ*^*^ _j_) and standard deviation (*σ*^*^ _j_) of log load for a genotype *j* (Eq. 2a, b):

<u>Eq. 2a: Equilibrium mean of log load</u>

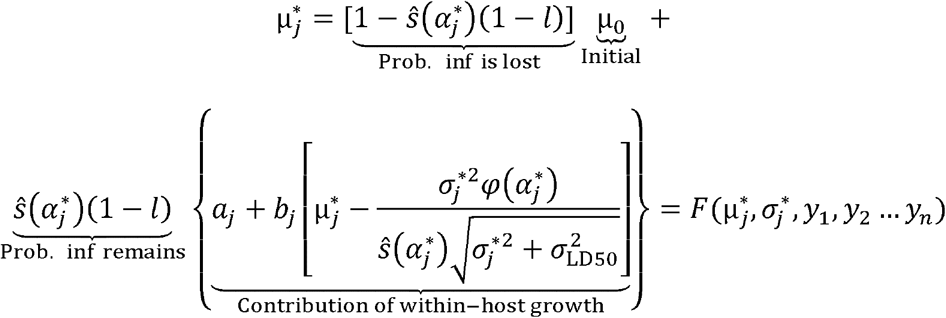

<u>Eq. 2b: Equilibrium variance plus mean squared of log load</u>

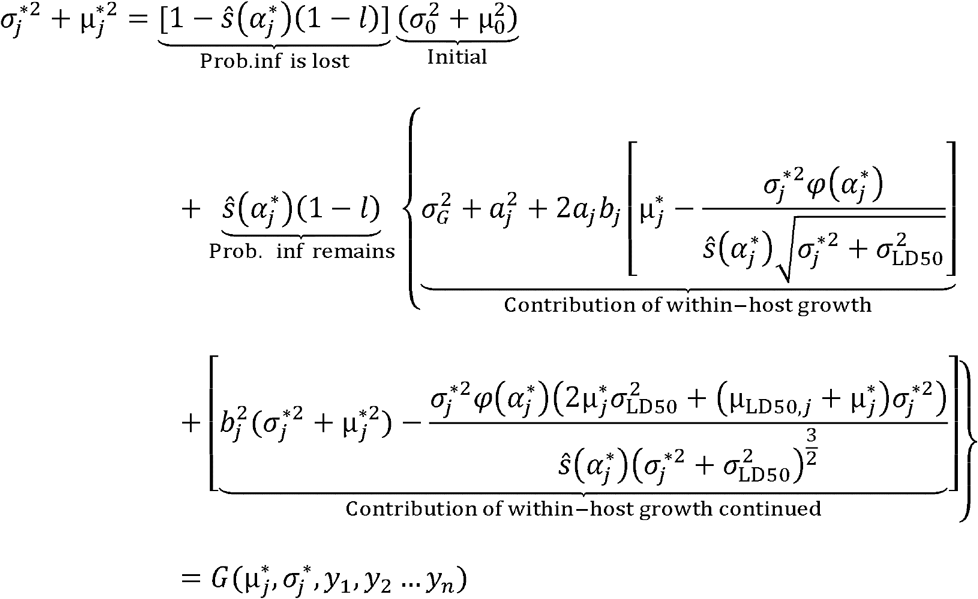

<u>Eq. 2c: Pathways affecting equilibrium mean log load</u>

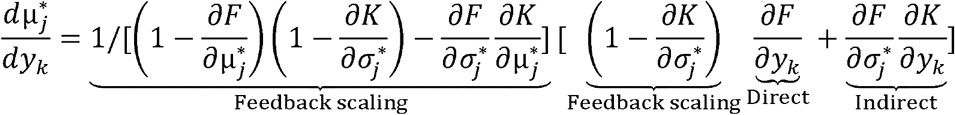

<u>Eq. 2d: Pathways affecting equilibrium SD of log load</u>

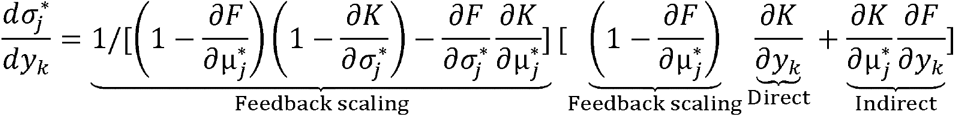

where *K* is the positive square root of *G*-*μ*_*j*_^*2^. The expressions for the mean and standard deviation of log load reflect the balance between the influences of new infection, pathogen growth on the host, and infection loss (particularly through death because it is load-dependent). The mean is the sum of initial mean weighted by the probability of infection loss plus the contribution of within-host growth (see terms in Eq. 2a that mirror terms in Eq. 1d) weighted by the probability that infection remains (i.e., an infected individual survives and does not recover). The expression for the equilibrium value of the variance plus mean squared follows the same structure (Eq. 2b). If we assume no loss of infection, we find explicitly that *µ*_j_^*^ = *a*/(1-*b*) and *σ*_j_^*^ = *σ*_G_/√(1-*b*). Because there is loss of infection, we retain the somewhat more complex expression in Eqs. 2a and 2b.

What factors influence the mean and standard deviation at equilibrium? We analyse these expressions to find how any parameter (*y*_*k*_ with *k* in 1 to *n*) affects *μ*^*^_j_ and *σ*^*^_j_, considered as functions of that fixed parameter. As long as the terms we label “feedback scaling” remain positive (Eqs. 2c, d; they do remain positive across a broad parameter range, see “Quantifying numerical robustness” in the Appendix), we can interpret these expressions as how *y*_*k*_ affects *μ*^*^ _j_ directly (“direct” term in Eq. 2c) and indirectly by affecting *σ*^*^ _j_ (“indirect” term in Eq. 2c). The direct term, ∂*F*/∂*y*_k_, captures how a change in the parameter *y*_k_ directly changes the *F* function (where *F* =*μ*^*^_j_). The indirect term comes from how *y*_k_ directly alters *σ*^*^_j_ (i.e., ∂*K*/∂*y*_k_) multiplied by how a change in *σ*^*^_j_ feeds back to change *μ*^*^_j_ (∂*F*/∂*σ*^*^_j_). The same logic applies for direct and indirect effects on *σ*^*^_j_ in Eq. 2d.

We find that pathogen growth and host death at high loads creates negative feedbacks between *μ*^*^_j_ and *σ*^*^_j_ (i.e., indirect effects, ∂*F*/∂*σ*^*^_j_, ∂*K*/∂*μ*^*^_j_, are negative). These negative feedbacks hold as long as certain inequalities, such as *µ*^*^ _j_ > *µ, σ*^*^ _j_^2^ + *µ*^*^_j_^2^ > *σ*_0_ ^2^+*µ*_0_^2^, and *µ*^*^_j_ < *µ*_LD50, j_ are met (see Appendix for one more inequalities and proofs; there is a fourth inequality that is sufficient but not necessary and less interpretable). Most of these assumptions are biologically reasonable, interpretable, and likely to hold for many systems. If *µ*^*^_j_ > *µ*_0_, then the loss of infections will bring the mean down as the lost infections are replaced with infections that have lower loads, on average. Similarly, if *σ*^*^_j_^2^ + *µ*^*^_j_^2^ > *σ*_0_^2^+*µ*_0_^2^, then the loss of infections will bring the variance plus mean squared down. If *µ*^*^_j_ < *µ*_LD50,j_ then fewer than 50% of infected hosts die each time step and increased variance increases death rate instead of decreasing it (see comment above about the effect of load variance on average survival). These negative feedbacks held across a broad range of parameter sets we numerically searched to compare to our results for our focal parameter set (see Quantifying Numerical Robustness section in the Appendix) and only failed to hold in parameter sets where *σ*^*^_j_^2^ + *µ*^*^_j_^2^ < *σ* _0_^2^+*µ*_0_^2^, *µ*^*^_j_ <*µ*_0_, or both (3.7% of parameter sets considered; these parameter sets seem unlikely to be broadly, biologically relevant). In summary, an increase in *µ*^*^_j_ increases the minority of infected hosts with high, deadly loads and their death squeezes the distribution of load, reducing *σ*^*^ _j_. An increase in *σ*^*^ _j_ also increases the minority of infected hosts with high, deadly loads and their death pulls *µ*^*^_j_ down. These mutual negative feedbacks arise because of growth of infection and the fact that the minority of hosts at equilibrium have lethal loads.

Understanding these mutual negative feedbacks allows us to understand how a perturbation shifts the distribution of load directly vs. indirectly by shifting these feedbacks. Consider only the *direct effects* of a perturbation that directly increases *µ*^*^_*j*_, shifting the distribution of load higher and having no direct impact on *σ*^*^_*j*_ ; biologically, this could be envisioned as a perturbation that changes all natural loads by the same percentage (because a fixed, multiplicative change in natural load is a fixed, additive change in log load), e.g., brief, favorable pathogen growth conditions that suddenly boost all pathogen loads by the same proportion. Every percentile of the load distribution is now found at higher load values, increasing disease-induced death rates for higher percentiles in particular. Then we consider the response in loads as enough time passes for the *indirect effects* of the perturbation to manifest. These higher percentiles of load experience higher death rates, shrinking the right tail of the distribution of load, *indirectly* decreasing *σ*^*^_j_ (see how the orange bell curve in Fig. 2D is narrower than the yellow bell curve). Similarly, a perturbation that only *directly* increases *σ*^*^_j_, say an increase in environmental stochasticity that increases stochasticity in pathogen growth (*σ*_G_), moves the left tail of load lower, with negligible impact on survival, and the right tail of load higher against the survival constraint, *indirectly* decreasing *μ*^*^_j_. Thus, a parameter that shifts both *μ*^*^_j_ and *σ*^*^_j_ also acts via their feedbacks; e.g., increased recovery rate (*l*) directly decreases *μ*^*^_j_ and *σ*^*^_*j*_ because both are constrained to be closer to initial infection levels; but lowering *μ*^*^_*j*_ strongly reduces the survival-based constraints on *σ*^*^_*j*_, creating a net increase in *σ*^*^_*j*_ ; this example helps us understand how various parameters affect *μ*^*^_*j*_ and *σ*^*^_*j*_ directly and via their mutual negative feedbacks (Table 1).

**Table 1.** Analytical results for direct and numerical results for total effects of parameters on *μ*^*^_j_ and _(_σ^*^_j_. Superscripts in columns 2 and 3 denote conditions analytically found to be sufficient (though not always necessary) for that sign to hold (see Appendix for derivations); the feedback scaling terms are always positive in a broad, biologically relevant parameter range. Columns 2 and 3 analytically determine the signs of the direct effects of parameter *y*_k_ on *μ*^*^_j_ and *σ*^*^_j_ respectively. Columns 4 (d*μ*^*^_j_ /d*y*_k_) and 5 (d*σ*^*^_*j*_ /d*y*_k_) quantify the total effects (direct effects+indirect effects, each multiplied by relevant feedback scaling from Eq. 2) found for the wild-type genotype in our focal parameter set. Mutual negative feedbacks between *μ*^*^_j_ and *σ*^*^_*j*_ guarantee that indirect effect of any *y*_k_ on *μ*^*^_j_ always has the opposite sign as *y*_k_’s direct effect on *σ*^*^_j_, and vice versa, due to mutual negative feedbacks. Parameters not listed do not affect *μ*^*^_j_ and *σ*^*^_j_ (e.g., note that *β*_j_ does not appear in Eqs. 2a, b).

| Changing parameter | Direct effect<br>on mean:<br>$\partial F / \partial y_k$ | Direct effect<br>on SD:<br>$\partial K / \partial y_k$ | Total effect<br>on mean<br>$d\mu_j^* / dy_k$ | Total effect<br>on SD<br>$d\sigma_j^* / dy_k$ |
| --- | --- | --- | --- | --- |
| SD of load upon initial<br>infection: $\sigma_0$ | 0 | $\geq 0$ | -0.073 | 0.078 |
| Mean load upon initial<br>infection: $\mu_0$ | $\geq 0$ | $\geq 0^1$ | 0.20 | -0.094 |
| $a_j$ contributes to pathogen<br>growth rate at all loads | $\geq 0$ | $\geq 0^{2, 3}$ | 5.1 | 0.80 |
| $b_j$ contributes to pathogen<br>growth rate at high loads | $\geq 0^{2, 3}$ | $\geq 0^{2, 3, 4, 5}$ | 26 | 16 |
| $\sigma_{LD50}$ makes the survival<br>function more gradual | $\leq 0^{2, 4, 6}$ | $\leq 0^{2, 5, 6, 7}$ | -0.64 | 0.020 |
| $\mu_{LD50,j}$ is the LD50 of the survival<br>function | $\geq 0^{2, 6}$ | $\geq 0^{2, 4, 8}$ | 0.42 | 0.042 |
| $\sigma_G$ controls stochasticity in pathogen growth | 0 | $\geq 0$ | -1.5 | 1.6 |
| Recovery rate from infection $l$ | $\leq 0^6$ | $\leq 0^8$ | -28 | 6.8 |

<sup>1</sup> $\mu_0 \geq 0$ , i.e., newly infected hosts have non-negative mean log load.
<sup>2</sup> $\mu_j^* < \mu_{LD50,j}$ , i.e., infected host survival probability for one day is $> 50\%$ .
<sup>3</sup> $\mu_j^* > 0.8\sigma_j^*$ , i.e., SD is not too much larger than mean.
<sup>4</sup> $\alpha_j^* < -1$ , i.e., infected host survival probability for one day is $> 84.1\%$ .
<sup>5</sup> $\mu_j^* > 0.177\mu_{LD50,j}$ , i.e., mean is not too much lower than LD50.
<sup>6</sup> $\mu_j^* > \mu_0$ , i.e., infection grows so that mean log load is greater than the mean of new infections
<sup>7</sup> $\alpha_j^* < -(3^{1/2})$ , i.e., satisfied if infected host survival probability for one day is $> 95.8\%$
<sup>8</sup> $\mu_j^{*2} + \sigma_j^{*2} > \mu_0^2 + \sigma_0^2$ . Guaranteed if condition 6 is met and $\sigma_j^* > \sigma_0 + 40$

These direct and indirect effects give us broadly applicable, conceptual insight into how various processes will impact the distribution of pathogen load. The signs of direct effects of these various parameters on *μ*^*^_*j*_ and *σ*^*^_*j*_ can be analytically guaranteed given biologically meaningful, sufficient conditions (see conditions under Table 1 and proofs in Appendix). Many of these conditions depend on the equilibrium, average probability of infected hosts surviving for one day, *ŝ*(*α*_*j*_^*^), not being too low; the most restrictive of these sufficient conditions is *ŝ*(*α*_j_^*^) > 0.958, which may be biologically reasonable given that we find *ŝ*(*α*_j_^*^) = 0.984 for our focal parameter values (see Table A1 for biologically reasonable parameter values and sources) and even higher for genotypes that evolve defence (also see Quantifying Numerical Robustness for the percentage of parameter sets in which these conditions held). Thus, we believe the signs of the direct effects that we found will hold in many biologically meaningful scenarios, giving us broad, conceptual insight into load dynamics. Additionally, these findings help us see how various processes would impact load, e.g., lower *a*_j_ due to higher constitutive resistance should strongly decrease mean and have a weak impact on the standard deviation; a population with a higher variance in initial load (*σ*_0_), e.g., due to some greater host variability, should have a higher standard deviation of load due to the direct effects of that heterogeneity but lower mean due to feedback and indirect effects. Further, feedbacks, direct effects, and indirect effects may prove insightful for the equilibrium distributions of traits other than load.

### Load signals in evolutionary recovery

The mutual negative feedbacks between the mean and standard deviation of load greatly aid our ability to infer differences in underlying mechanisms via differences in load. Due to their mutual negative feedbacks, the mean and standard deviation of load can shift in opposite directions in response to a change in a single mechanism (the direct effect columns of Table 1 never have opposite sign but the net effect columns do). This greatly improves our ability to infer underlying processes from the distribution of load as the mean and standard deviation can each have three qualitative responses (+, flat, -) and nine theoretical combinations of responses. We demonstrate the utility of these load signatures with a focal case study of Mountain Yellow Legged Frog populations recovering from *Bd*.

Population abundance and genetic data indicate Mountain Yellow Legged frog populations are recovering in the presence of a deadly fungal pathogen (*Bd*) due to some form of evolution relevant to host defences [25]. However, the precise process of host evolution is unknown. Frogs may evolve tolerance that improves their survival of high load infections (higher *μ*_LD50, j_ in our model), resistance that prevents infection (lower *β*_j_), or resistance that decreases pathogen growth. Resistance to pathogen growth may be constitutive and apply equally to all infections (lower *a*_j_), be inducible to disproportionately resist growth of high load infections (lower *b*_j_), or be acquired immunity that only protects reinfected frogs (lower *a*_j_ in reinfected frogs; see Appendix for the extension of Eq. 1 to the case of acquired immunity). These different processes correspond to different timing and likely molecular mechanisms of host immunity evolution. We simulated outbreak of fungal disease in a population of mostly wild-type frogs with a low starting frequency of nine other genotypes spanning a range of defence in one of these traits with a fecundity cost of this higher defence (see Appendix). The outbreak of the pathogen drives host density down (Fig. 3A) as prevalence increases (Fig. 3B) but the more defended genotypes then increase in frequency (Fig. 3C) and alter the distribution of load (Fig. 3D and E) with implications for pathogen abundance (Fig. 3F).

**Figure 3.**
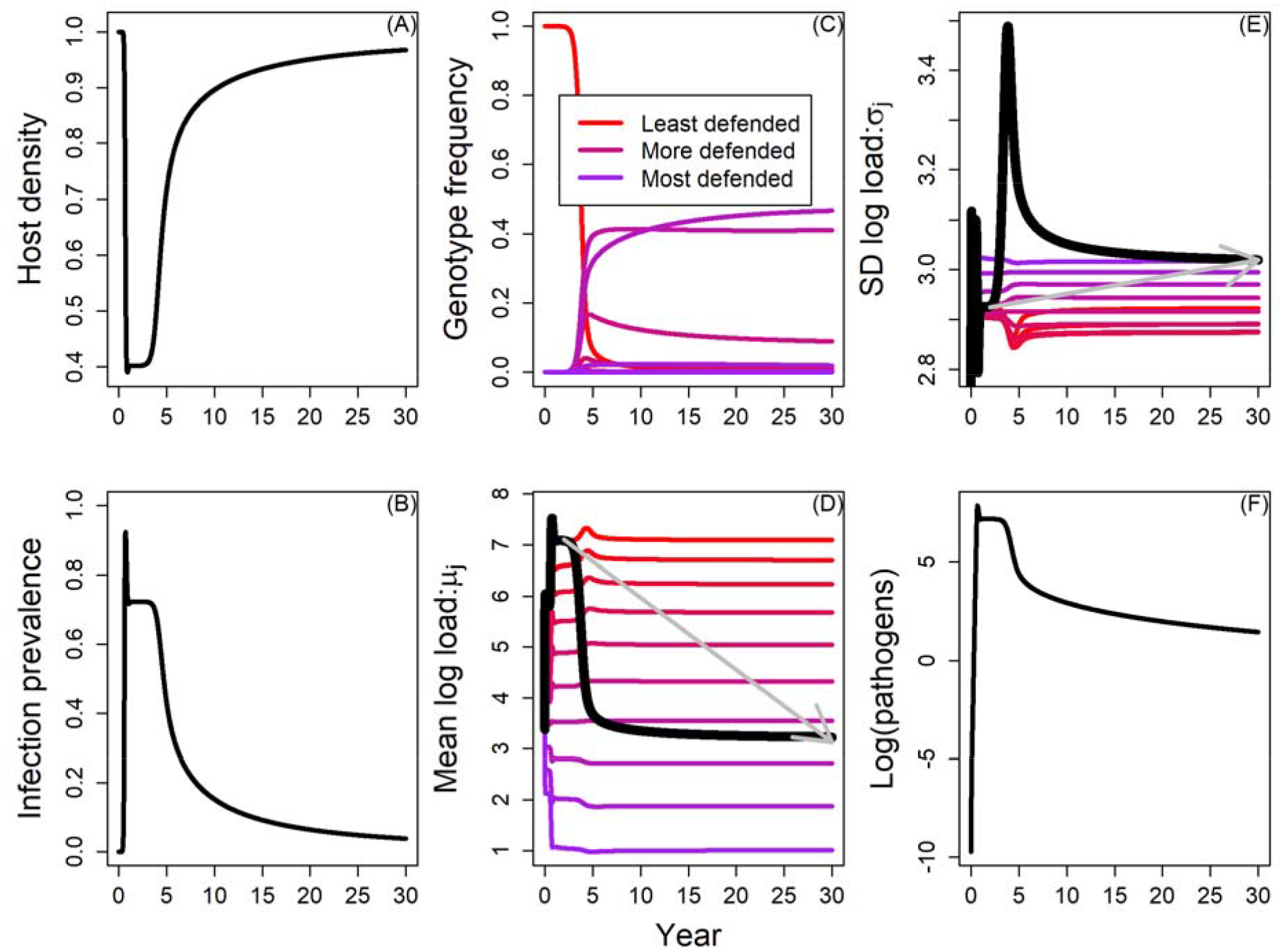
Evolutionary recovery via constitutive resistance. Host genotypes vary in the *a*_*j*_ parameter and more defended genotypes (lower *a*_*j*_) have lower fecundity. (A) Pathogen invasion causes host density to decline then recover as hosts evolve. (B) Invasion increases prevalence but then evolution of constitutive resistance decreases prevalence. (C) Host evolution occurs as the frequencies of clonal genotypes shift from the least defended, wild-type genotype to more defended genotypes. (D) More defended genotypes have lower mean log loads (coloured curves) and the overall mean log load of the entire infected population declines as hosts evolve resistance (black curve with gray arrow emphasizing change from wild-type stable equilibrium until the end of the simulation). (E) The standard deviation of load has a weaker, more mixed dependence on constitutive resistance but weakly increases overall as hosts evolve. Note a transient spike in the overall standard deviation around year 5 due to a temporary mixture of more and less well defended genotypes. (F) Host evolution decreases pathogen density in the environment.

Critically, we find that these different defence hypotheses correspond to qualitatively different patterns in pathogen load. As more defended genotypes increase in frequency, the overall load pattern in the entire infected host population generally follows that predicted by the equilibrium analysis for a single genotype in Table 1 (though Table 1, alone, is insufficient to predict the reinfection case). Increasing tolerance allows a higher and wider spread of loads, leading to a higher mean and weakly higher standard deviation (see Fig. A1 in Appendix). Resistance that prevents infection has no direct impact on the mean and standard deviation but changes the timing of epidemic stage, leading to weak and transient changes in the mean and standard deviation (see Fig. A3). Different forms of resistance to pathogen growth (depicted graphically in Fig. 4A) also correspond to qualitatively different trends. Constitutive resistance decreases the mean and weakly increases the standard deviation (Fig. 4B); this result arises because constitutive resistance has a weak, negative, direct effect on the standard deviation but the strong decrease in mean and negative feedbacks of the mean on the standard deviation leads to a net positive effect on the standard deviation (see the row for *a* in Table 1). Inducible resistance has a strong, direct, negative impact on both the mean and standard deviation (Fig. 4C). Acquired resistance to pathogen growth in reinfections drives lower mean and strongly higher standard deviation (Fig. 4D); this increase arises because the infected population is now a mixture of hosts infected for the first time, with relatively high loads, and reinfected hosts, with lower loads (see Appendix for explanation of when and why this increases the standard deviation). This increases the variance in log load in the total infected host population. These results were highly robust to variation in the model parameters and arose in the model without the simplifying assumption of lognormal load (see Quantifying Numerical Robustness section in the Appendix), though the model without the simplifying assumption provides less conceptual clarity as to the reason for these results than our focal model here does. Thus, the distribution of load can qualitatively differentiate among hypotheses for the defence evolution of Mountain Yellow Legged Frogs, or other hosts.

**Figure 4.**
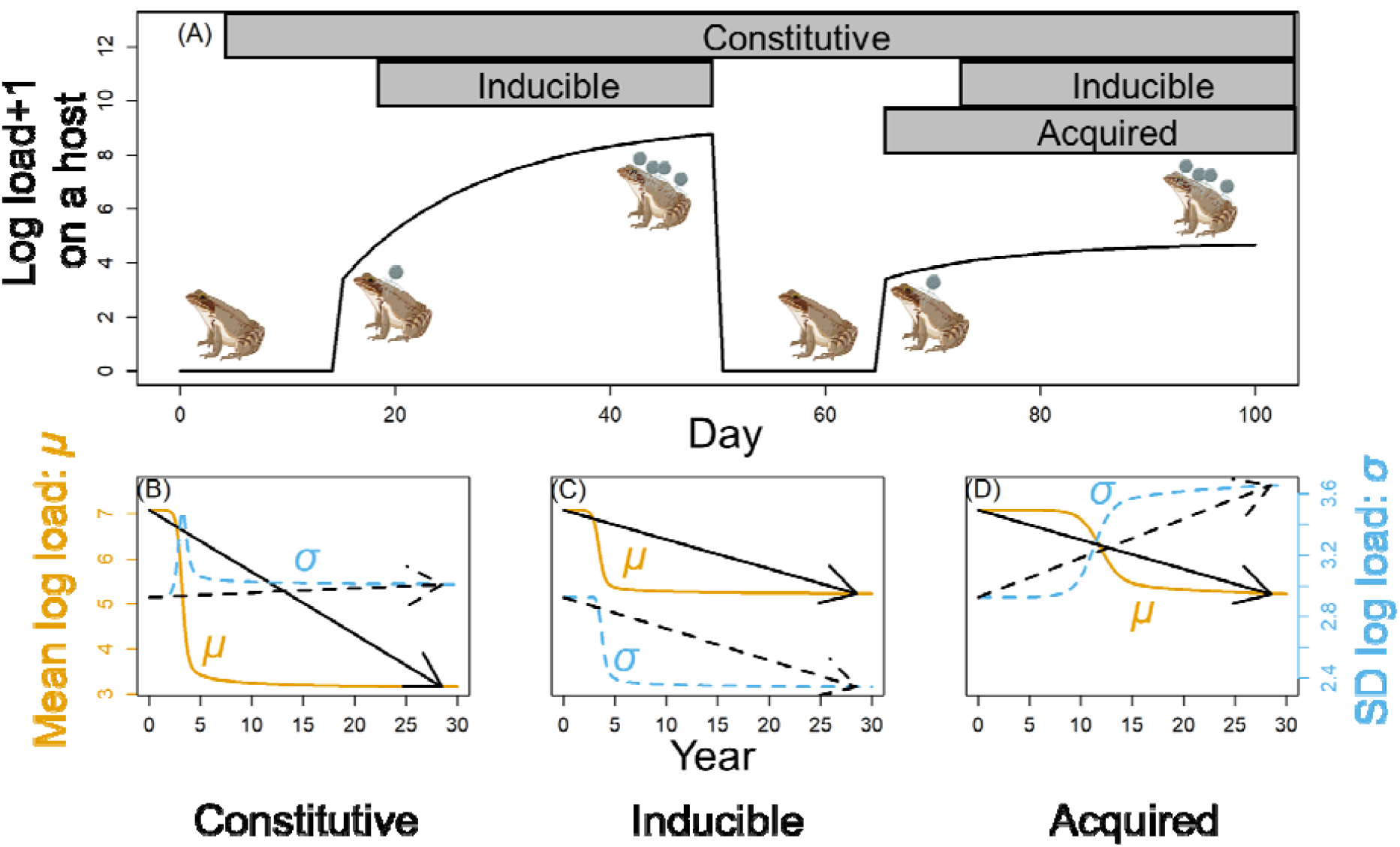
Load trends qualitatively differ for different forms of resistance. (A) We show deterministic ln(load+1) on a single host witsh all three types of resistance tracked over 100 days with infection on day 16, recovery on day 51, and reinfection on day 66. Constitutive resistance is always on. Inducible resistance acts most strongly at high loads. Acquired resistance only applies to reinfections. (B) Increasing constitutive resistance decreases mean log load (gold) while the standard deviation increases weakly (blue). Black arrows emphasize the overall trends. (C) Inducible resistance decreases the mean and strongly decreases the standard deviation. (D) Acquired resistance decreases the mean and increases the standard deviation because the entire infected population is now composed of first-time infected hosts with high mean loads and reinfections with lower mean loads. These simulations were initialized at the wild-types endemic equilibrium with low frequencies of the nine, more defended genotypes (see Appendix).

### Impacts of the load distribution on population dynamics

We exemplify the impact of the distribution of load on population dynamics by contrasting how different forms of resistance affect host and pathogen fitness as well as endemic conditions. Consider the dynamics of a single genotype with strong constitutive resistance experiencing an epidemic in isolation contrasted to those of a single genotype with strong inducible resistance. Both genotypes keep mean loads low but constitutive resistance has a higher standard deviation of log load at equilibrium (*σ*^*^_Constitutive_ > *σ*^*^ _Inducible_); because mortality falls disproportionately on high load infections (assuming *μ*^*^_Constitutive_ < *µ*_LD50_ and *μ*^*^ _Inducible_ < *µ*_LD50_, i.e., fewer than 50% of infected hosts die each day), the higher standard deviation of constitutive resistance contributes to higher death rates. For constitutive and inducible resistance to achieve the same, average fitness while infected [*ŝ*(*α*^*^ _Constitutive_) = *ŝ*(*α*^*^ _Inducible_)], the genotype with constitutive resistance must reduce mean log load further [we set *a* = 0.35 for the constitutive genotype and *b* = 0.88 for inducible with all other parameters as wild-type; see Fig. 5A; *µ*\*_Constitutive_ < *µ*\*_Inducible_ is required by *σ*\*_Constitutive_ >, *σ*\*_Constitutive_ *μ*\*_Inducible_ < *µ* _LD50_, and *σ*\*_Inducible_ < *µ*_LD50_]. Despite lower mean loads, the higher standard deviation of the constitutively resistant genotype means there are more individuals with rare, very high loads that lead to higher average pathogen shedding rates per infected hosts [*µ*\*_Constitutive_ +*σ*^*^_Constitutive_^2^/2 > *µ*^*^_Inducible_ +*σ*^*^_Inducible_^2^/2; see Appendix for biologically reasonable conditions under which this inequality will hold]. Higher pathogen shedding rates contribute directly to pathogen fitness (see *R*_0_ expression in [23]) and host population depression in the case of constitutive resistance compared to inducible resistance (Fig. 5B). Therefore, different forms of host defence and thus standard deviations of load can have strongly different implications for pathogen fitness, even given equal host fitness while infected (acquired resistance falls on this same continuum, see Fig. A6).

**Figure 5.**
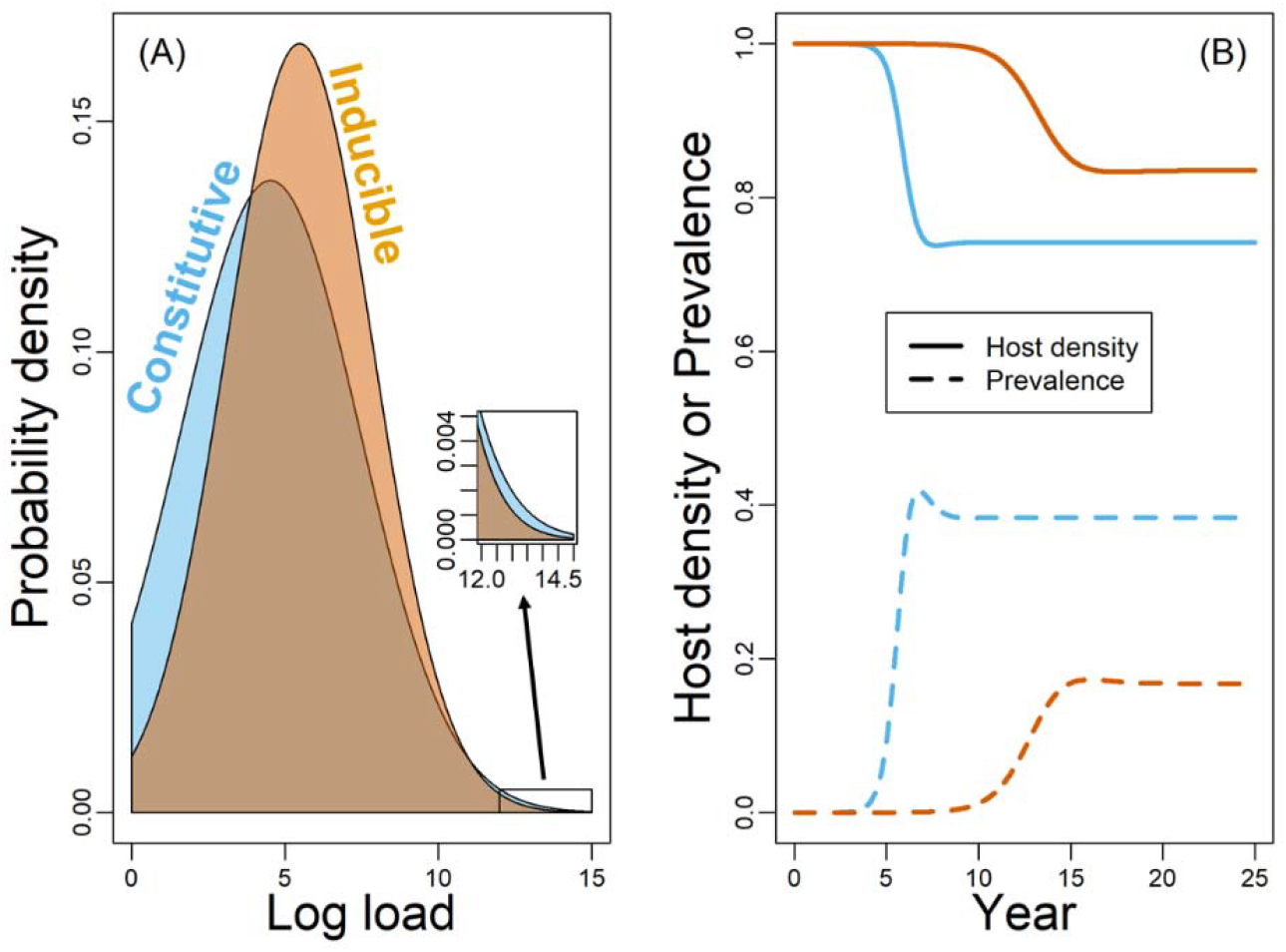
Higher standard deviation contributes to host death, pathogen spread, and host population depression. We show results for single-genotype simulations for a genotype that either possesses strong constitutive (*a* = 0.35; all other parameters as wild-type) or inducible (*b* = 0.88) resistance. For comparability, we parameterized these to give the same, average infected host fitness at equilibrium. (A) Constitutive resistance (blue) leads to a lower mean and higher standard deviation of log load compared to inducible resistance (orange). Constitutive resistance then has a slightly higher proportion of hosts with very high loads (see magnified inset panel). (B) More high load infections lead to stronger host population depression for constitutive resistance than inducible (solid blue lower than solid orange) due to greater pathogen spread (dashed blue higher than dashed orange).

## Discussion

How does variance in microparasite load interact with population-level dynamics? We show how epidemic stage alters the mean and standard deviation of load due to the rate of influx of new infections, pathogen growth, and high mortality at high loads. Further, we show that the mean and standard deviation have negative feedbacks with each other due to high mortality at high loads. These negative feedbacks create qualitatively different trends in the mean and standard deviation for different forms of host defence evolution, improving our ability to infer within-host processes from population-level patterns. Finally, we show how different trends in the variance in load, e.g., due to different forms of host defence evolution, have different impacts on host fitness, pathogen fitness, and host suppression by pathogens. Thus, we have gained new, conceptual insights into the interactions among variance in microparasite load and population-level dynamics that improves our understanding of the focal frog-*Bd* system and likely many other microparasite systems.

Many of our key results will apply to microparasite load broadly. While other microparasites may differ structurally in their between-host processes (e.g., frequency-dependent transmission instead of environmental transmission), many of our key findings depend primarily on assumptions regarding within-host processes that should be more general. Microparasite infections grow beyond their initial load, negative density-dependence within hosts constrains the loads that can be attained, survival probability declines in some non-linear fashion from one to zero as loads become very high, and the minority of infected hosts dies in a given time step.

Thus, many of our key, conceptual findings should apply to other microparasites despite differences in parameter values or details of model structure. For example, our finding that mean loads rise as an epidemic slows, due to a reduced influx of new infections bringing the mean down, should apply to many microparasites. Negative feedbacks between the mean and standard deviation, due to a non-linear survival constraint, should also arise frequently as they depend on conceptually reasonable conditions, such as mean loads being higher than initial loads and less than half of infected hosts dying every day; these negative feedbacks should allow different forms of host defence to generate qualitatively different impacts on mean and standard deviation in other microparasite systems as well. Further, our specific model and findings will be particularly, directly applicable to microparasites that display lognormal distributions of load like HIV and SARS-CoV-2 [13,14,20,21]. Because, any of our key results were supported analytically and those supported numerically had broad numerical support, we conclude that our key results should apply broadly to many other microparasite systems.

We were able to draw broader conclusions because our model was relatively simple and tractable for analytical work, showing broad conditions in which our patterns would hold. The relative simplicity of our model relies on a lognormal distribution of pathogen load, which is empirically supported for *Bd* [24] and likely many other microparasites [13,14,20,21]. Second, our model provides insights that are not readily available in simpler models with dynamic mean and fixed variance [23] or in more complex models that make no simplifying assumptions about the distribution of load [22]. Without our simplifying assumption of lognormal load, it would have been much more difficult to derive expressions for the mean and variance of load at equilibrium and clarify the existence of and reason for various key results. For mutual negative feedbacks, for example, our simplifications allowed us to find that the decline in host survival at high loads leads to negative feedbacks between the mean and the standard deviation. Thus, any change (e.g., a change in a host trait) that directly affects mean and standard deviation also indirectly affects them by shifting their feedbacks (Table 1). Disentangling these direct and indirect pathways clarifies how the distribution of log load changes given a parameter perturbation, e.g., constitutive resistance (lower *a*) directly lowers the standard deviation but can actually cause it to weakly increase due to the sum of direct and indirect effects (Figs. 3 and 4). A notable result from our modelling approach is that different forms of resistance have qualitatively different impacts on the variance of load (Fig. 4), greatly improving our ability to infer within-host processes from population-level patterns as well as predict population-level impacts of epidemics (Fig. 5).

Improved understanding of load dynamics, in general, also helps us recognize how the stochastic processes governing the standard deviation of load can become a battleground for evolution by hosts and/or microparasites. Our model confirms the predictions from classic compartmental models that tolerance (higher *µ*_LD50_ in our model) increases prevalence and environmental pathogens (Råberg et al. 2008), confirming the clear intuition that tolerance should increase mean log loads but illustrates with greater novelty that tolerance also increases the standard deviation of log load. Resistance mechanisms, on the other hand, should decrease pathogen fitness [26], as we found. We were able to differentiate resistance mechanisms even further. We found that different forms of resistance had different impacts on the variance in load (Fig. 4). Higher standard deviation, e.g., associated with constitutive resistance vs. inducible resistance, increases the likelihood of exceptionally high load infections, leading to host death. Even assuming lower mean log load for constitutive resistance to achieve the same average death rate of infected hosts, we found that higher variance should lead to higher pathogen shedding per infected host, increasing pathogen fitness, prevalence, and depressing the host population more for constitutive resistance than inducible. Thus on the spectrum of resistance (increasing host fitness and decreasing pathogen) to tolerance (increasing host fitness without decreasing pathogen fitness), constitutive resistance is more “tolerant” than inducible resistance. Previous compartmental models of constitutive vs. inducible resistance have considered how the two forms of resistance may differ in whether their costs and benefits are expressed while susceptible or infected [27,28]; but we assume that all three of our forms of resistance, constitutive, inducible, and acquired, have constant costs and only provide benefits while infected yet we still find they have different implications for the standard deviation, host fitness, pathogen fitness, pathogen prevalence, and host density. Thus, dynamic heterogeneity in pathogen load reveals important differences among different forms of resistance and points the way toward new considerations for host-parasite coevolution.

Our work builds toward a greater understanding of how load dynamics shape host and microparasite fitness (and thus perhaps coevolution) at the within- and between-host scales. Previously, nested models have demonstrated that within-host dynamics, like pathogen growth, influence tradeoffs between transmission rate and virulence, impacting pathogen evolution at different epidemic stages [29]. And a previous moment closure of an IPM with dynamic mean and constant variance showed that faster within-host growth can increase pathogen fitness through higher shedding or decrease it if host mortality becomes too high [23]. Our work emphasizes the importance of a dynamic standard deviation of load for both host and pathogen fitness for their impact on pathogen spread as well as our ability to infer underlying processes.

These results suggest new avenues for studying host-pathogen coevolution. For example, evolution by host or microparasite will affect how strongly stochastic mechanisms can generate heterogeneity in load, altering the standard deviation that controls the probability of very high load infections that shed at high rates (good for pathogens) and die quickly (bad for hosts). Further, a model like ours could help investigate more fully the likely critical feedbacks between the within-host and between-host scales, a criterion that makes such models particularly vital [30]. For example, high force of infection from the environment may increase the mean load of a new infection (would create a *β*_j_*Z* dependence of our *G*_0_ function and allow avoidance resistance to alter equilibrium load dynamics). Further, pathogen growth rates may depend critically on dose (as dose has been found to influence many within-host dynamics: Kamiya et al. 2020) such that low dose infections are eliminated by hosts but high dose infections grow out of control (we could readily model this by modifying our *a*+*bx* growth parameters to have *a* < 0, *b* > 1 or using some non-linear function with a third parameter). Further empirical work could determine the relevance of such functional forms while further theoretical work could determine their implications, e.g., possible bistability between endemic and disease-free equilibria.

In addition to multiple conceptual advances, our model also provides a methodological advance for certain applications to merge theory and data. We demonstrate that investigators may be able to use the load dynamics of the mean *and* standard deviation to infer underlying processes, such as which form of defence hosts are evolving. Additionally, the reduced dimension of our model with the moment closure, compared to without the moment closure, will improve the efficiency of fitting a model to population-level data (similar to the moment closure in Bolker and Pacala 1997 for improving fitting efficiency for a different problem) including host abundance, infection prevalence, and load dynamics. Because our model links within-host and population-level processes, such a fitting procedure may be very useful for combining multiple data types. For example, laboratory data may provide informative prior probability distributions for some parameters, improving the ability of a model fitting procedure to infer other within-host processes from population-level data. In our focal system, Mountain yellow-legged frogs are likely evolving some form of resistance to pathogen growth [18,25,33]; ample data are collected on abundance, infection prevalence, and the distribution of *Bd* load in recovering, wild populations so that fitting a version of our model to these data may identify whether this resistance is more likely constitutive, inducible, or acquired helping target future immunological investigations and conservation actions (e.g., whether or not to immune prime reintroduced frogs). While we use temporal trends in load patterns to infer temporal differences in within-host processes, the same approach could be used for other trends (e.g., latitudinal trends in load and within-host processes). Our model also serves as a resource for investigators interested in lognormally distributed pathogens and may inspire investigators to use this same moment closure approach to derive helpful models for other distributions (e.g., gamma distributions).

Further, our findings call for more investigation of how dynamic trait heterogeneity can inform relatively simple ecological models for new insight. Our approach could be applied directly to track dynamic mean and variance of other lognormally distributed, ecological traits outside of disease systems (e.g., body size), and is particularly useful when there are non-linear transition probabilities (e.g., survival), e.g., to Stubberud et al. 2019 [34]. Some of our insights would apply most readily to traits that tend to change directionally over time, like body size, so that the loss of individuals constrains the mean and variance closer to starting values, depending on mean-variance feedbacks (see Table 1 for how these can work out). Our assumption of a normal distribution of a non-heritable trait parallels the common simplifying assumption in quantitative genetics of a normal distribution of a heritable, heterogeneous trait [1]. Additionally, we found that trends in the variance of load can be very informative regarding underlying mechanisms, a concept that may apply well to quantitative genetics. For example, [2] used a quantitative genetics model to show how changes in mean trait can reveal upcoming population shifts; within their model framework, changes in the trait variance may also reveal mechanisms by which the mean shifts as we expect a change via selection should decrease trait variance (at least temporarily) while a shift via adaptive plasticity may not. Given the conceptual insights that can be obtained with relatively simple models, we recommend further investigations of how heritable and non-heritable sources of trait variation interact to shape trait variance and population-level processes.

Overall, our model results emphasize the value of considering load dynamics for critical, ongoing questions of microparasite diseases. Previous theory and theory-data integration have effectively utilized the mean of infection load, providing a strong theoretical foundation and many datasets to leverage when considering a dynamic standard deviation of infection load. Relatively simple models like ours provide computationally efficient, empirically relevant, and analytically tractable tools for advancing our understanding of load dynamics for microparasites. Further, we can address how classic aggregation metrics change with these models to see how microparasite dynamics follow, or deviate from, patterns found for macroparasites. Improving our understanding of load dynamics will allow us to infer within-host processes more efficiently, predict host population resilience more accurately, and build a more complete understanding of (co-)evolution of host-microparasite systems.

## Notes

### Competing Interest Statement

The authors have declared no competing interest.

### Summary of Updates

We have revised the text and figures to emphasize the biological conclusions and broader implications of this work more fully.

### Funding

- National Science Foundation: DBI-2120084

